# Hackflex library preparation enables low-cost metagenomic profiling

**DOI:** 10.1101/2024.04.23.590092

**Authors:** Samantha L. Goldman, Jon G. Sanders, Daniel D. Sprockett, Abigail Landers, Weiwei Yan, Andrew H. Moeller

## Abstract

Shotgun metagenomic sequencing provides valuable insights into microbial communities, but the high cost of library preparation with standard kits and protocols is a barrier for many. New methods such as Hackflex use diluted commercially available reagents to greatly reduce library preparation costs. However, these methods have not been systematically validated for metagenomic sequencing. Here, we evaluate Hackflex performance by sequencing metagenomic libraries from known mock communities as well as mouse fecal samples prepared by Hackflex, Illumina DNA Prep, and Illumina TruSeq methods. Hackflex successfully recovered all members of the Zymo mock community, performing best for samples with DNA concentrations <1 ng/uL. Furthermore, Hackflex was able to delineate microbiota of individual inbred mice from the same breeding stock at the same mouse facility, and statistical modeling indicated that mouse ID explained a greater fraction of the variance in metagenomic composition than did library preparation method. These results show that Hackflex is suitable for generating inventories of bacterial communities through metagenomic sequencing.

## Introduction

The ability to sequence the entire complement of DNA present in an environmental sample (i.e., metagenomic shotgun sequencing) has rapidly expanded knowledge of the structure and function of microbial communities. However, metagenomic shotgun sequencing remains cost prohibitive for many studies, prompting development of methods that reduce the cost of metagenomic DNA library preparation [1–3]. One of these methods, Hackflex, adapts the Illumina DNA Prep protocol (formerly labeled Nextera Flex) by diluting bead-linked transposases during tagmentation and using commercially available reagents yielding libraries at 1/14th the cost [3].

Hackflex has been widely validated and used in the context of whole-genome sequencing for bacterial isolates [4–11] and amplicon sequencing [12]. Although Hackflex has been used in metagenomic analyses, its accuracy has only been tested in the context of simple mock communities containing seven species of known relative abundances [13, 14]. Thus, the utility of Hackflex for metagenomic shotgun sequencing of complex microbial communities from natural sources has not been robustly evaluated.

Here, we tested the effectiveness of Hackflex for metagenomic sequencing by assessing its ability to accurately recover bacterial DNA from a known mock community and to profile bacterial communities in mouse fecal samples. We compare Hackflex to the higher-cost Illumina DNA Prep and TruSeq library preparation methods. We show that Hackflex libraries accurately recover known Zymo mock communities and identify template DNA concentrations that minimize observed biases in relative abundance estimates. When applied to mouse fecal DNA, we show that Hackflex libraries recapitulated results obtained from costlier Illumina DNA Prep and TruSeq libraries. Moreover, Hackflex was sufficiently sensitive to differentiate the metagenomes of individual mice from the same inbred line reared in a common environment. These results support the utility of Hackflex for metagenomic studies.

## Results and Discussion

### Hackflex recapitulates Zymo mock communities

To assess the accuracy of metagenomes sequenced from libraries prepared by Hackflex (Supplementary Methods), we sequenced Hackflex libraries prepared from the ZymoBIOMICS Microbial Community DNA Standard (Catalog # D6306) —a cocktail of DNA from 8 bacterial and 2 fungal species mixed at known proportions—using a range of template DNA concentrations. In total, 18 Zymo mock DNA communities were sequenced at seven different DNA template concentrations generated by serial 1:2 dilutions of each sample (Table S3; Supplemental Methods). Details about library fragment size and sequencing depth are presented in Supplementary Results.

To assess the accuracy of Hackflex, we next used a custom kraken2 [15] database containing genomes from all Zymo mock community members to measure the difference between the observed and known relative abundance for each community member. The percentages of reads that mapped to the custom database with minimap2 [16] ranged from 96.85% to 98.34% (mean = 97.93%) (Table S3), consistent with minimal contamination in these experiments. Results indicated that Hackflex successfully replicated the expected relative abundances of the bacterial (12%) and fungal (2%) components of the Zymo mock DNA community within one order of magnitude (Fig 1A, 1B), although an overrepresentation of *Lactobacillus* and an underrepresentation of *Listeria, Bacillus, Enterococcus, Escherichia, Salmonella*, and *Pseudomonas* was observed (Fig 1B).

**Figure 1.**
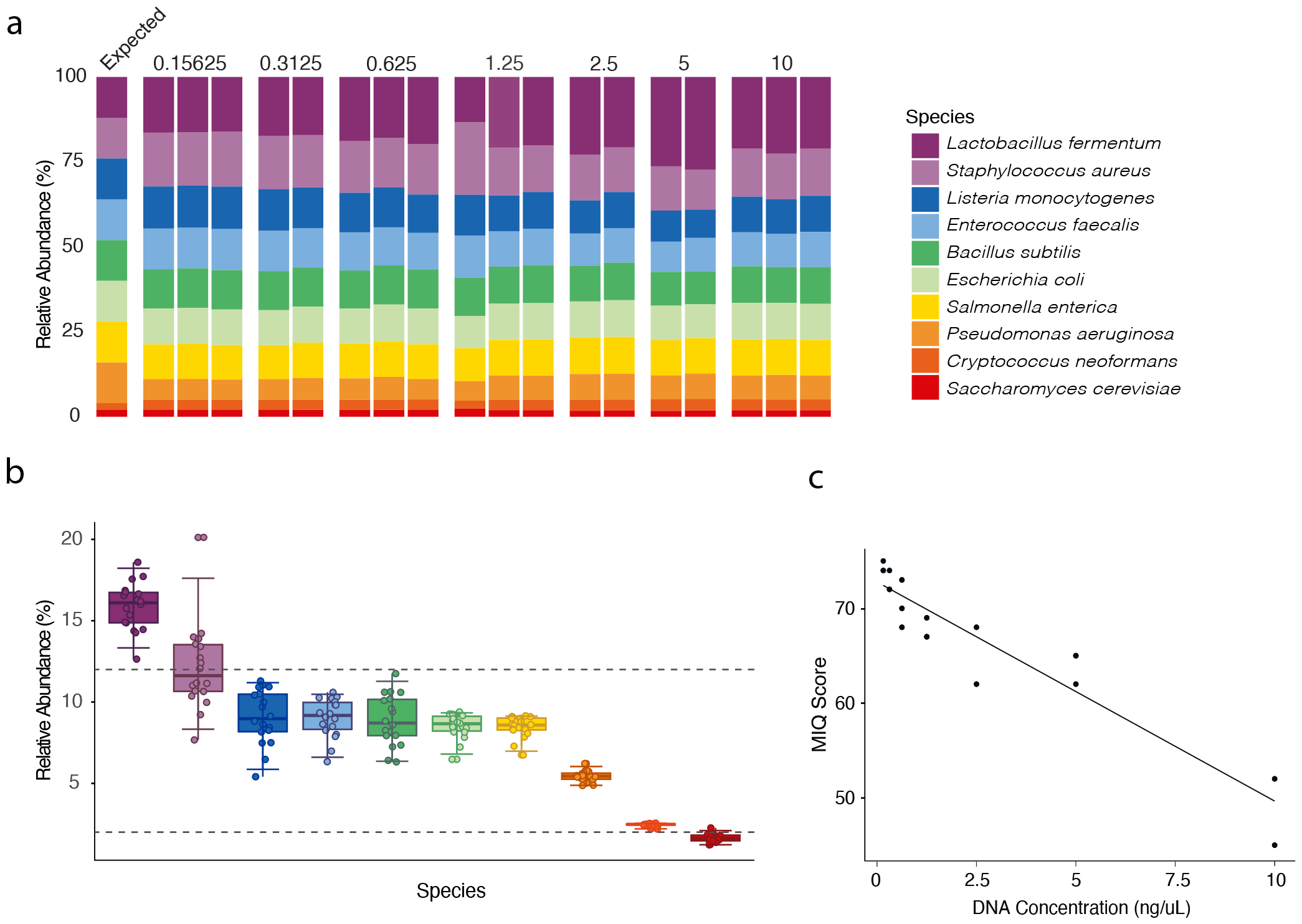
Hackflex recovers mock community metagenomes. A) Normalized taxa barplots show expected and observed relative abundance for mock communities at each DNA concentration (ng/uL). Colors denote microbial species as indicated by the key. Leftmost barplot shows expected relative abundances. B) Boxplots show observed relative abundances for each taxon in each sample. Colors correspond to those in (A). Points represent relative abundances of microbial species. 95% confidence intervals are represented in pink. Horizontal lines show expected relative abundances for bacteria (top line) or eukaryotes (bottom line). C) Scatterplot shows MIQ Scores and DNA concentrations for each sample. Points represent libraries generated from Zymo mock DNA. Line indicates best-fit regression.

In addition, for each Zymo metagenome we calculated Measurement Integrity Quotient (MIQ) scores, which assess bias by calculating deviations from the expected relative abundance of each taxon while allowing for variance inherent in the manufacturing of mock communities [16]. Results from these analyses indicated that Hackflex yielded passing scores (i.e., >60) for 16/18 samples, with only libraries prepared from DNA samples at highest concentration (10 ng/uL) failing (mean = 66.778, median = 68) (Fig 1C, Table S3, Supplementary Fig 1). We additionally calculated the MIQ scores for samples from a previous study which prepared five Illumina DNA Prep libraries from the Zymo mock DNA community [17] (mean MIQ score = 86.8, median = 86) (Supplementary Table 3). These results show that, at low input DNA concentrations, Hackflex yields MIQ scores ∼86% as high as those obtained from Illumina DNA Prep, while allowing savings of over 10-fold lower reagent costs.

We observed no significant biases as a function of microbial domain, gram status, GC content, or genome size (Table S2, Supplementary Fig 3). However, Hackflex MIQ scores were significantly negatively associated with DNA concentration (R^2^ = 0.876; p = 1.19e-8) (Fig 1C, Table S3). One explanation for this association is that decreasing the ratio of beads to DNA shifts the tagmentation reaction such that DNA molecules compete more for DNA tagmentase when DNA concentrations are high, exacerbating any underlying biases. These results indicate that Hackflex’s biases can be reduced by diluting template DNA to ∼0.15 ng/uL.

### Hackflex corroborates Illumina DNA Prep and TruSeq when applied to mouse gut metagenomes

We also tested the accuracy of Hackflex for biological samples (mouse fecal metagenomes) by comparing its performance with Illumina DNA Prep (Fig 2). We prepared 25 total libraries from five fecal samples from five different mice. To prioritize technical replication for Hackflex libraries, we prepared four serial 1:2 dilutions per Hackflex-prepared sample (Fig S1, Table S1). Details about library fragment size and sequencing depth are presented in Supplementary Results. For downstream analyses, Illumina DNA Prep samples were used as a reference against which Hackflex samples were compared for accuracy.

**Figure 2.**
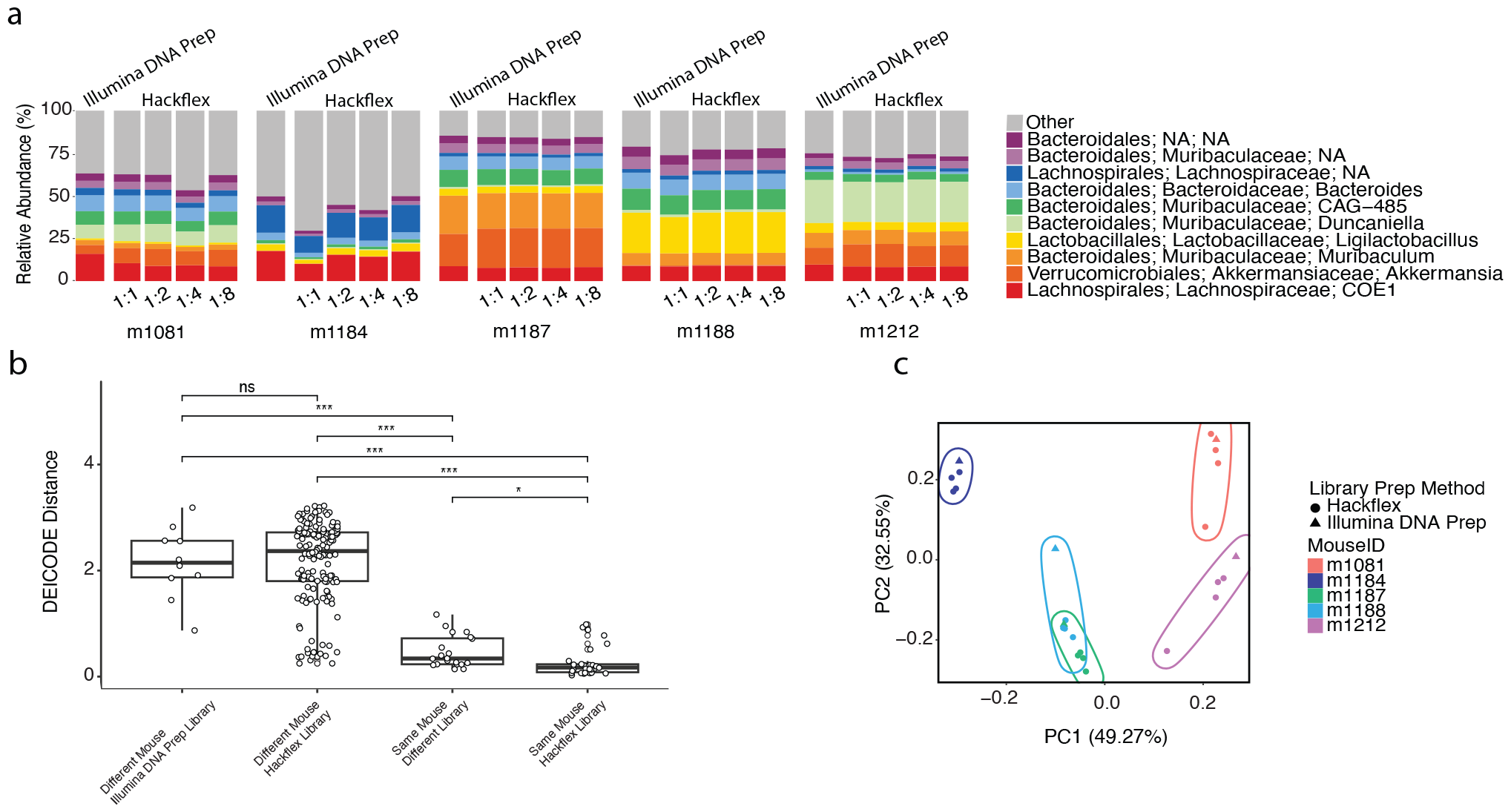
Hackflex recovers individual signatures in mouse gut metagenomes. A) Taxa barplots show relative abundances of microbial taxa observed in mouse metagenomes sequenced from libraries prepared with Illumina DNA Prep or Hackflex. Colors denote microbial genera as indicated by the key. B) Boxplots show the DEICODE Aitchison dissimilarities between pairs of samples from different mice prepared with Illumina DNA Prep, different mice prepared with Hackflex, the same mice prepared with different library methods, and the same mouse prepared with Hackflex. Points represent pairwise comparisons between samples. Asterisks and ‘ns’ indicate significance of differences between boxplots based on permutation t-tests for non-independent samples; Bonferroni-corrected *p*-value < 0.05 *; < 0.001 ***; > 0.05 ns. C) Principal coordinate analysis plots show Robust Aitchison dissimilarities among samples. Circles represent Hackflex-prepared libraries and triangles represent Illumina DNA Prep-prepared libraries. Colors denote individual mice as indicated by the key.

Hackflex performed comparably to Illumina DNA Prep in recovering metagenomes from mouse fecal samples (Fig 2A). PERMANOVA based on Woltka [19] taxonomic profiles generated for all samples indicated that mouse ID was a stronger driver of community variation (R^2^ = 0.222) compared to library prep method (Illumina DNA Prep vs Hackflex) (R^2^ = 0.053) (Table S2). Pairwise robust Aitchison DEICODE [20] comparisons between samples showed that taxonomic profiles of samples collected from the same mouse but whose libraries were prepared by different methods were more compositionally similar than those of libraries collected from different mice but whose libraries were prepared by the same method (Fig 2B) (*p*-value < 0.001; pairwise permutation t-test for non-independent samples, Bonferroni multiple comparison correction). Furthermore, principal coordinate analysis of DEICODE [20] distances showed that these samples clustered by mouse individual rather than library preparation method (Fig 2C). The two mice that were the least distinguishable, m1187 and m1188, were cagemates and therefore expected to share many microbes [21] (Table S4). In addition, we repeated tests of the accuracy of Hackflex by comparing results obtained from a different set of five mouse fecal samples for which libraries were prepared with both Hackflex and Illumina TruSeq (Supplementary Results, Figure S4), revealing qualitatively identical results to the comparisons between Hackflex and Illumina DNA Prep.

This study shows that Hackflex represents an effective, low-cost library preparation method for shotgun metagenomics. Biases of the method may skew estimates of bacterial relative abundances, but these biases can be mitigated by dilution of template DNA to concentrations as low as ∼0.15 ng/ul.

Moreover, Hackflex was able to recover individual-host signatures in the mouse gut microbiota, even among mice of the same genotype reared in a common environment. Therefore, Hackflex reduces costs for inventorying microbial taxa and estimating their relative abundances within complex microbial communities.

## Data Availability

Raw sequences uploaded to NCBI archive under accession number XXXX. The R code used to generate figures is available at https://github.com/samanthagoldman/hackflex_validation/ and at the Zenodo record https://zenodo.org/doi/10.5281/zenodo.10965899.

## Acknowledgements

We thank Hannah Grazul for assistance with DNA extractions and Hackflex library preparations from rodent fecal samples.

