## Supplemental Methods for "Hackflex library preparation enables low-cost metagenomic profiling"

*Design of experiments*

To evaluate the biases of the Hackflex library preparation method, we generated and sequenced libraries from DNA from a mock microbial community. These analyses used the ZymoBIOMICS Microbial Community DNA Standard (D6306), a mixture of DNA from eight bacteria and two yeasts. The abundance of each taxon is scaled for genome size so that the DNA concentration is equal for each group. Genomic DNA from each bacterial and yeast species is at 12% relative abundance and at 2% relative abundance, respectively.

While the mock community is ideal for systematically validating Hackflex, it does not represent the complexities of a full gut microbial community. Therefore, we additionally tested fecal samples from C57BL/6 mouse (*Mus musculus domesticus*) individuals (Supplementary Tables 1, 4) ordered from the Jackson Laboratory (https://www.jax.org). Mice were reared under standard laboratory conditions and then a subset were transferred to an outdoor enclosure for 10 days prior to fecal sampling. All animal work was performed under Cornell University International Animal Care and Use Committee protocol number 2015-0060.

*Library preparation, pooling, sequencing, and processing*

DNA from mouse fecal samples was extracted and Hackflex libraries from both the mouse samples and mock DNA community were prepared as described in Sanders et al. [4]. In short, DNA was extracted from fecal samples using a magnetic bead-based protocol on an OpenTrons OT2 liquid handling robot (https://github.com/CUMoellerLab/Moeller_Opentrons_protocol_library/tree/master/Library_Prep/Hackflex). Libraries for both the mouse samples and mock DNA community were prepared using Hackflex library preparation as previously described [4]. TruSeq and Illumina DNA Prep libraries were prepared at the Cornell University Biotechnology Resource Center. TruSeq-Hackflex comparison libraries and Zymo mock DNA community libraries were sequenced on an Illumina NextSeq 500, and Illumina DNA Prep-Hackflex comparison libraries were sequenced on an Illumina NovaSeqX as previously described [4]. For each sequencing run (Illumina NextSeq 500 or Illumina NovaSeqX), pools of samples were multiplexed in equimolar amounts and first sequenced on an Illumina MiSeq Nano Kit v2. Data from the MiSeq Nano sequencing run was then used to infer, based on read counts, the relative abundances of each sample in the original pools. These sample DNA relative abundances were then used to re-pool DNA from each sample to equal amounts, as previously described [4], before resequencing on Illumina NextSeq 500 or Illumina NovaSeqX.

*Bioinformatic analyses of mock community samples*

Raw sequencing reads were imported into Qiita [22]. Within Qiita, adapters and host-derived (mouse) sequences were removed from the mouse gut samples using qp-fastp minimap2 (v2021.01). Processed reads were imported into QIIME 2 (v2023.9) [23]. To determine whether Hackflex recovered taxonomic groups present in the Zymo mock DNA community, we created a custom database of genomes present in the mock DNA community using Struo2 (v2.3.0) [24] and mapped reads derived from these samples using kraken2 (v2.3.1) [15]. The number and percentage of unclassified reads for each of these samples is presented in Supplementary Table 3. Taxa barplots were generated with the qiime2R package (v0.99.6) [25]. We utilized the Zymo-developed Measurement Integrity Quotient (MIQ) [17] to reproducibly determine whether Hackflex led to any biases within our dataset. MIQ scores were assigned by measuring the root mean square error of observed abundances compared to the known abundance of each taxon in the sample. MIQ scores can range from 0-100, with 100 indicating no bias and 0 indicating maximum bias.

Linear regression of DNA concentration and MIQ scores was performed using the lm function from base R (v4.3.2) [26]. Linear regression and ANOVA analyses to test for biases were conducted on center log ratio transformed data in base R. Correction for multiple testing was performed using p.adjust in base R [26]. Adjusted p-values for sensitivity analyses were calculated separately from those calculated from the whole dataset. The summary statistics for these analyses are reported in Supplementary Table 2.

*Bioinformatic analyses of mouse fecal samples*

Taxonomic profiles of mouse fecal samples were estimated with Woltka (v0.1.1) [17], a classifier that maps reads to the Web of Life, a broad microbial phylogenetic database [23]. Qiita-generated feature tables were imported into QIIME2 (v.2022.2) [24] and taxa bar plots were generated. DEICODE, a form of Aitchison distance that is robust to sparse datasets [20] was calculated on species-level taxonomic profiles using the qiime deicode rpca function (Fig 2B). PERMANOVA was conducted with the adonis2 function in the vegan package (v2.6.4) [28] (Supplementary Table 2).

To determine whether there was a bias in GC content or gram status, we performed taxonomic assignment with MetaPhlAn4 (v4.0.6) [29] for TruSeq vs. Hackflex samples. For any named species that was present in at least both TruSeq and Hackflex-prepared samples form a given mouse, we searched the name of the species on NCBI Taxonomy to obtain GC content for the reference genome for that species. We then searched species names on BacDive: The Bacteria Diversity Metadatabase (<https://bacdive.dsmz.de>) to obtain gram status. Accession numbers for the reference genomes and gram status are presented in Supplementary Table 5. Relative abundance estimates were center log ratio transformed to reduce effects of compositionality. Linear regression and ANOVA analyses to test for biases were conducted in base R. Correction for multiple testing was performed using p.adjust in base R [27]. The summary statistics for these analyses are reported in Supplementary Table 2.

**Supplemental Results**

*Library fragment size and read counts*

Library pools were checked for quality on a TapeStation using ProSize Data Analysis Software. For the Hackflex-versus-Illumina DNA Prep comparisons, the mean fragment size for Hackflex libraries was 236bp and the mean fragment size for Illumina DNA Prep libraries was 501bp. For the Hackflex-versus-Illumina TruSeq comparisons, the mean fragment size for Hackflex libraries was 241bp and the mean fragment size for Illumina TruSeq libraries was 527bp. For these comparisons, >95% of library fragments prepared by Hackflex were between 100bp and 500bp, whereas >95% of library fragments prepared by Illumina DNA Prep or TruSeq were between 250bp and 1000bp, as expected. The final concentration of the pools sequenced were 1.402 ng/uL and 1.016 ng/uL for the Hackflex-versus-Illumina TruSeq comparison and Hackflex-versus-Illumina DNA Prep comparison, respectively.

In the final Zymo mock DNA community dataset, each sample yielded on average 1,207,773 reads (standard error = 59,383) before adapter and quality filtering and 926,423 reads (standard error = 40,947) after quality filtering. For the TruSeq v. Hackflex comparison experiment, each TruSeq library yielded on average 5,141,263 reads (standard error = 889,054) before adapter, host-read, and quality filtering and 3,715,651 reads (standard error = 1,001,029) after filtering. One Hackflex library sample was removed from further analyses as it yielded only one read. The remaining Hackflex-prepared samples yielded an average of 1,314,593 reads (standard error = 51,575) per sample before adapter and quality filtering and 638,986 (standard error = 75,094) reads per sample after filtering. For the Illumina DNA Prep versus Hackflex comparison experiment, the Illumina DNA Prep libraries yielded on average 8,575,658 reads (standard error = 2,100,353) per sample before adapter, host-read, and quality filtering and 7,652,590 reads (standard error = 1,897,295) after filtering. The Hackflex libraries yielded on average 4,143,779 reads (standard error = 276,978) per sample before adapter, host-read, and quality filtering and 2,311,362 reads (standard error = 236,831) after filtering. The lower coverage for Hackflex libraries relative to TruSeq or Illumina DNA Prep libraries enabled additional technical replication for Hackflex libraries. In both comparisons, <50,000 reads were observed for the negative control samples (i.e., blank wells containing no template DNA but for which library preparation was conducted), consistent with minimal contamination in our library preparations.

*Hackflex was not biased as a function of microbial domain, gram status, GC content, or genome size*

To determine whether results from Hackflex libraries are biased as a function of domain (Bacteria vs Eukaryota), gram status (for bacterial taxa), GC content, or genome size, we performed ANOVAs (for domain and gram status) and linear regressions (for GC content and genome size) based on the deviation between centered log ratio transformed relative abundances of each Zymo species with each of these traits (Supplementary Figure 2). These analyses revealed no significant effects of domain, gram status, GC content, or genome size after accounting for multiple testing (Supplementary Table 2). A modest effect of genome size was observed when only bacteria were considered, but sensitivity analyses showed that the significance of this result (at the p-value <0.05 level) was dependent on a single species (*L. fermentum*). Together, these results fail to identify any biases of data obtained by Hackflex library preparation method as a function of domain, gram status, GC content, or genome size.

*Hackflex corroborates TruSeq when applied to mouse gut metagenomes*

We also tested the accuracy of Hackflex for biological samples (mouse fecal metagenomes) by comparing its performance with that of TruSeq, a widely used and more costly method for metagenomic sequencing. We prepared 15 total libraries from five fecal samples from five different mice, with between two and four 1:2 dilutions per Hackflex-prepared sample (Supplementary Fig 4A, Supplementary Table 1). We used a homemade TruSeq equivalent method that uses the same ligase as the commercial Illumina TruSeq library prep [30]. For downstream analyses, TruSeq samples were used as references against which Hackflex samples were compared for accuracy. Given the higher library costs, we sequenced TruSeq libraries at approximately 4x the depth of Hackflex and prioritized technical replication for Hackflex libraries.

Hackflex performed comparably to TruSeq in recovering metagenomes from mouse fecal samples (Supplementary Fig 4A). PERMANOVA based on Woltka [19] Web of Life [23] taxonomic profiles generated for all samples indicated that mouse ID was a stronger driver of community variation (R^2^ = 0.309) compared to library prep method (TruSeq vs. Hackflex) (R^2^ = 0.151) (Supplementary Table 2). However, in one mouse fecal sample dominated by Lactobacillus (m1082), Hackflex failed to recover the remaining diversity in this sample. This result is consistent with the results from Zymo mock DNA communities that indicated a bias towards Lactobacillus. Pairwise DEICODE comparisons between samples showed that taxonomic profiles of samples collected from the same mouse but whose libraries were prepared by different methods were more compositionally similar than were those of samples collected from different mice but whose libraries were prepared by the same method (Fig. 2B) (p < 0.001, permutation t-test for non-independent samples, Bonferroni multiple comparison correction). Furthermore, principal coordinate analysis of robust Aitchison distances using DEICODE [20] showed that these samples clustered by mouse individual rather than library preparation method (Supplementary Fig 4C). Interestingly, the two mice that were the least distinguishable, m1192 and m1194, were cagemates and therefore expected to share many microbes [21] (Supplementary Table 4). We did not observe any bias with regard to GC content (Bonferroni-corrected *p-*value = 0.197) (Supplementary Fig 3) or gram status (Bonferroni-corrected *p*-value = 1) (Supplementary Table 4).

**Supplementary Figure and Table Captions**

**Supplementary Figure 1.** MIQ score diagrams for each mock DNA community sample. Rows represent technical replicates of the concentration of DNA. The black circle in each diagram represents an unbiased community. Values that lie outside the black circle are overrepresented and values that lie within the black circle are underrepresented. Microbes are sorted around each circle by GC content.

**Supplementary Figure 2**. Hackflex shows no bias with regard to GC content. Linear regression of mock DNA community samples sorted by GC content shows that Hackflex has no bias towards lower GC content (slope = -0.010856, t-statistic = -1.125, Bonferroni-corrected p-value p = 1). Red points are average difference for a given species.

**Supplementary Figure 3.** In a complex community, Hackflex shows no bias with regard to GC content in comparison to TruSeq. Linear regression of named species in mouse samples sorted by GC content shows that Hackflex is unbiased with regards to GC content in this community (slope = 0.008952, t-statistic = 0.643, Bonferroni-corrected p-value = 0.197). Red points are average difference for a given species.

**Supplementary Figure 4.** Hackflex recovers individual signatures in mouse gut metagenomes. A) Taxa barplots show relative abundances of microbial taxa observed in mouse metagenomes sequenced from libraries prepared with TruSeq or Hackflex. Colors denote microbial genera as indicated by the key. B) Boxplots show the DEICODE Aitchison dissimilarities between pairs of samples from different mice prepared with TruSeq, different mice prepared with Hackflex, the same mice prepared with different library methods, and the same mouse prepared with Hackflex. Points represent pairwise comparisons between samples. Asterisks and ‘ns’ indicate significance of differences between boxplots based on permutation t-tests for non-independent samples; Bonferroni-corrected *p*-value < 0.01 **; < 0.001 ***; > 0.05 ns. C) Principal coordinate analysis plots show Robust Aitchison dissimilarities among samples. Circles represent Hackflex-prepared libraries and triangles represent TruSeq-prepared libraries. Colors denote individual mice as indicated by the key.

**Supplementary Table 1. Sample metadata.**

**Supplementary Table 2. PERMANOVA formulae and tables.**

**Supplementary Table 3. Mock DNA Community MIQ Scores.**

**Supplementary Table 4. Mouse metadata.**

**Supplementary Table 5. Reference genomes used for bias analyses.**
