## Supplementary figures and images for "Hackflex library preparation enables low-cost metagenomic profiling"

### Supplementary Figure 1

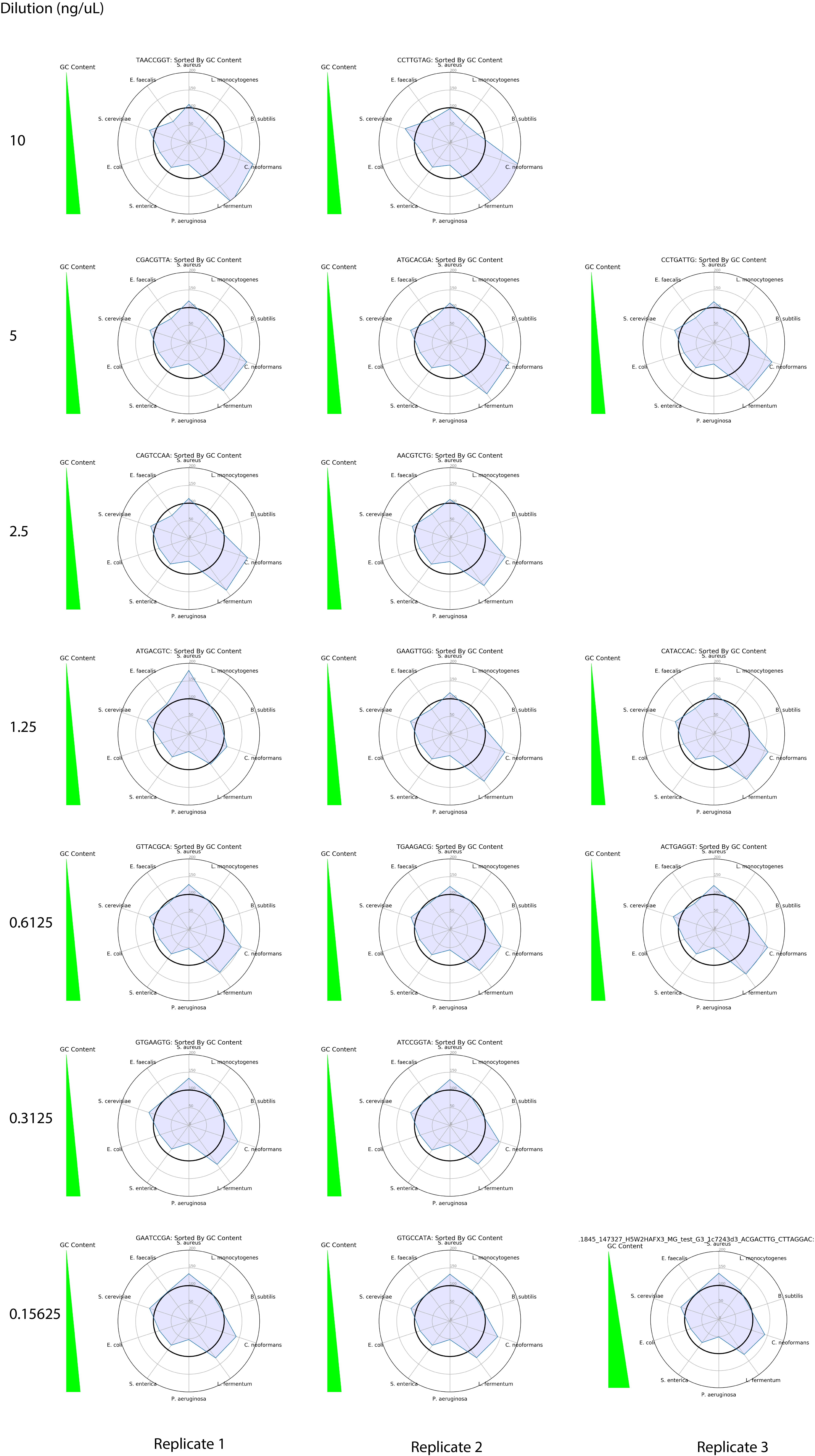

### Supplementary Figure 2

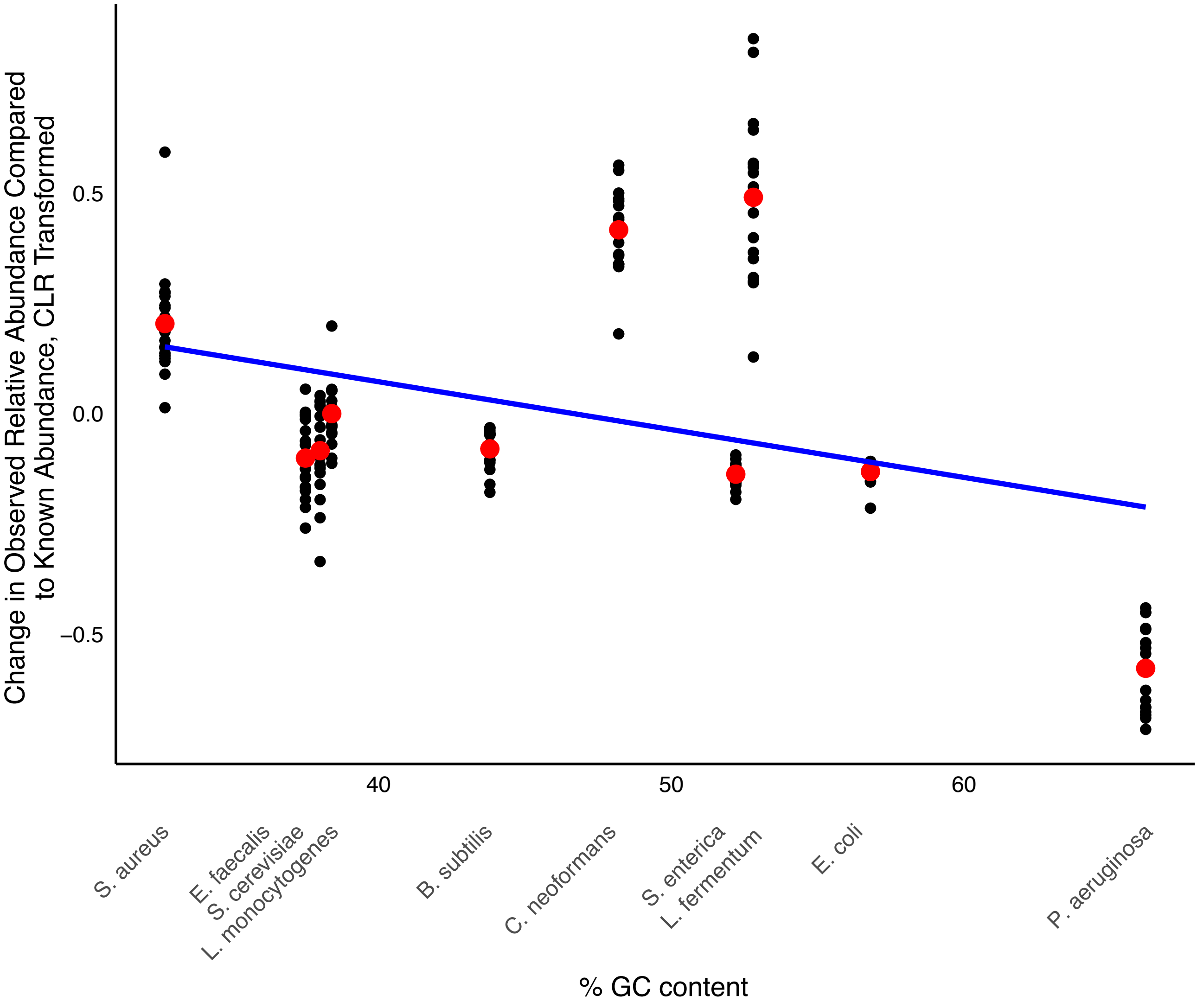

### Supplementary Figure 3

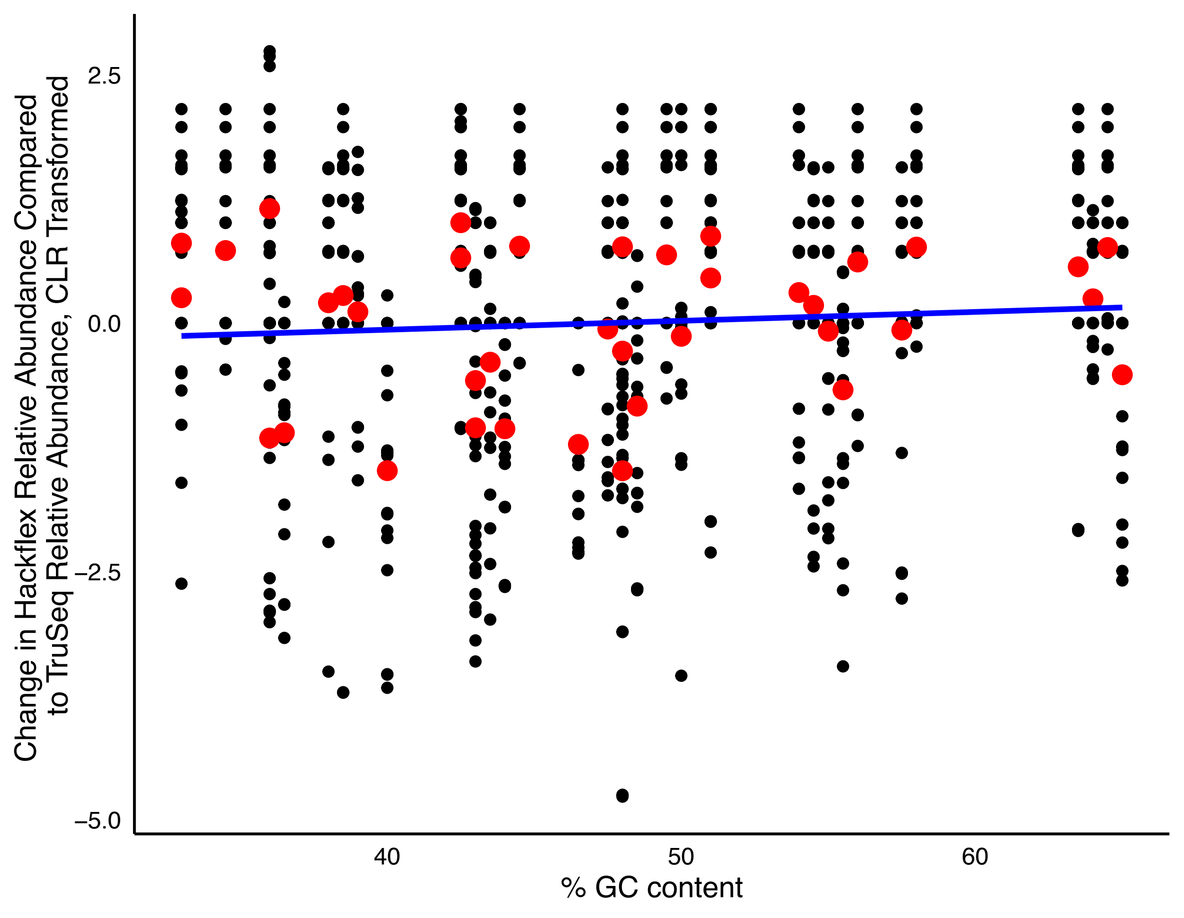

### Supplementary Figure 4

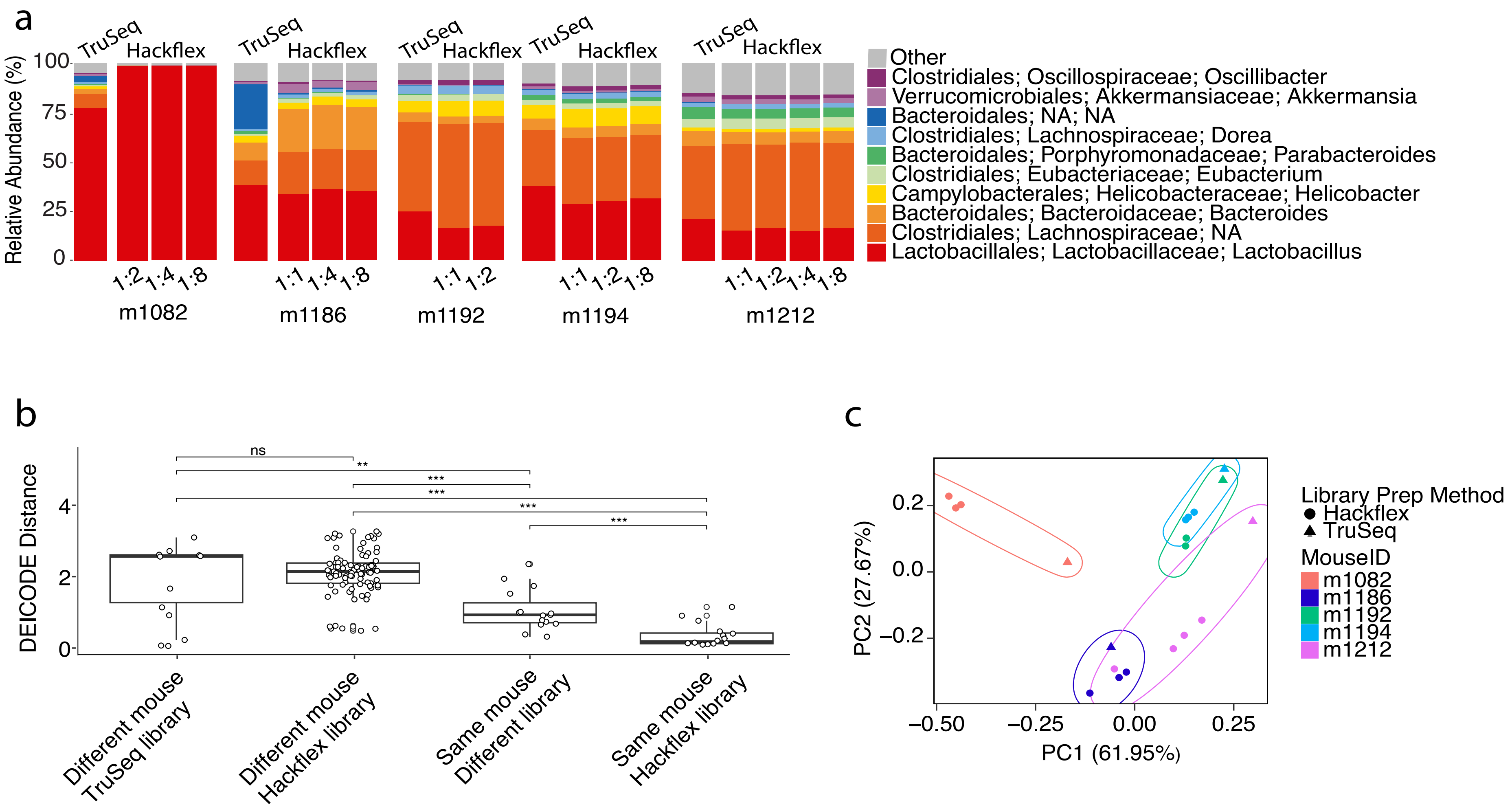
